# Individual differences in sequential decision-making

**DOI:** 10.1101/2025.04.04.647306

**Authors:** Mojtaba Abbaszadeh, Erica Ozanick, Noa Magen, David Darrow, Xinyuan Yan, Nicola Grissom, Alexander B. Herman, Becket R. Ebitz

## Abstract

People differ widely in how they make decisions in uncertain environments. While many studies leverage this variability to measure differences in specific cognitive processes and parameters, the key dimension(s) of individual variability in uncertain decision-making tasks has not been identified. Here, we analyzed behavioral data from 1001 participants performing a restless three-armed bandit task, where reward probabilities fluctuated unpredictably over time. Using a novel analytical approach that controlled for the stochasticity in this tasks, we identified a dominant nonlinear axis of individual variability. We found that this primary axis of variability was strongly and selectively correlated with the probability of exploration, as inferred by latent state modeling. This suggests that the major factor shaping individual differences in bandit task performance is the tendency to explore (versus exploit), rather than personality characteristics, reinforcement learning model parameters, or low-level strategies. Certain demographic characteristics also predicted variance along this principle axis: participants at the exploratory end tended to be younger than participants at the exploitative end, and self-identified men were overrepresented at both extremes. Together, these findings offer a principled framework for understanding individual differences in task behavior while highlighting the cognitive and demographic factors that shape individual differences in decision-making under uncertainty.

## 2 Introduction

Different people approach the same task in different ways. The way people respond in identical circumstances depends on their prior experiences, differences in cognitive styles and brain architectures, and on personal characteristics like age and gender. Understanding the latent factors that mediate these individual differences is critical for understanding the causes of individual variability, as well as for understanding and treating neurobiological and psychiatric conditions that affect task performance. However, in part because cognitive tasks are often complex, stochastic, and require the coordination of multiple cognitive processes, we do not always know which latent factors drive individual differences in performance. Developing methods to uncover these factors will be the key to identifying the cognitive processes(s) that a task is best suited to measure and to developing sensitive instruments for use in clinical populations.

One common and highly influential class of tasks in the decision-making literature is the k-armed bandit: a test of decision-making under uncertainty that manipulates the reward contingencies of a set of options (“arms”) that are evaluated in sequence. Individuals may approach bandit tasks differently as a function of sex (Chen et al. (2021a); Joue et al. (2022)), developmental stage (Kim and Carlson (2024); Xu et al. (2023); Sumner et al. (2019); Meder et al. (2021); Harms et al. (2024); Ohta et al. (2024); Mizell et al. (2024)), exposure to environmental stress (Kaske et al. (2023); Frankenhuis and Gopnik (2023); Xu et al. (2023)), personality characteristics (Yan et al. (2019); Jach et al. (2023); Lévy-Garboua et al. (2024); Dubois and Hauser (2022); Steyvers et al. (2009)), among many other factors. Critically, bandit tasks also have potential as a clinical tool (Knep et al. (2024); Chen and Vinogradov (2024); Dubois and Hauser (2022)). Bandit task performance differs with a variety of clinical diagnoses, including anxiety, depression, bipolar disorder, autism spectrum disorder, and psychosis (Chen and Vinogradov (2024); Aylward et al. (2019); Gilmour et al. (2024); Kaske et al. (2023); Knep et al. (2024); Addicott et al. (2013, Addicott et al. 2021); Harlé et al. (2015); Dubois and Hauser (2022); Thinzar (2024); Daumas et al. (2023)). As a result, understanding the latent factors that drive individual differences in bandit task performance could improve our understanding of sex, aging, chronic stress, and a variety of neurological conditions.

Many studies have leveraged bandit tasks to measure individual differences in specific cognitive factors. In particular, previous studies have focused on individual differences in reward processing (Zuo et al. (2025); Gilmour et al. (2024); Bouneffouf et al. (2017)), decision noise (Averbeck (2015); Yan et al. (2024); Findling and Wyart (2021); Wang et al. (2025)), cognitive flexibility (Lee et al. (2025); Skowron (2024)), decision-making strategies (Schulz et al. (2018); Lee et al. (2025); Chen et al. (2021a); Horvath et al. (2021)), and explore-exploit trade offs (Cohen et al. (2007); Averbeck (2015); Brown et al. (2022); Kim and Carlson (2024); Addicott et al. (2017)). While all of these latent cognitive factors are correlated with some aspect of bandit task performance or strategy, it remains unclear which of these factors (if any) are the primary driver of inter-individual variability in these tasks. The goal of this study is to identify the latent factors that account for individual variability in bandit task performance and determine whether these factors are linked to specific demographic characteristics.

In order to identify the latent cause of individual variability in bandit tasks, we developed a novel method for measuring the manifold of individual task strategies via a nonlinear and model-free embedding of task behavior. In a large dataset of people performing a classic k-armed bandit task, we find that the principle axis of inter-individual variability is strongly and selectively aligned with the probability of exploration, as identified by latent state models. In terms of demographics, we find that both gender and age predict variation between individuals along this axis: self-identified men are more common at the extremes of the manifold and older adults tend to be less exploratory than younger adults. Mathematical analyses revealed that the changes with age were primarily due to differences in the frequency or stability of exploratory behavior. Together, these results suggest that the k-armed bandit task is ideally suited to measure individual differences in exploration and highlight the utility of this task for understanding the effects of age on exploration, decision-making, and cognitive flexibility.

## 3 Results

A total of 1001 participants performed 300 trials each of an uncertain sequential decision-making task known as a probabilistic restless 3-armed bandit (Fig. 1A; see Methods 5.2). In this task, three options were associated with reward probabilities that changed randomly and unpredictably over time (Fig. 1B). Performance in uncertain decision-making tasks like this one is shaped by multiple cognitive factors. First, because participants can only infer action value by sampling options and integrating experienced rewards over time, this task is commonly used to estimate certain parameters of reinforcement learning processes (Barto (2021); Daw et al. (2006); Rescorla (1972); Zid et al. (2024)). Second, participants must choose how to allocate their options between known good options and alternative options that could become better at any time, making this task well suited for measuring individual differences in exploratory behavior (Wilson et al. (2021); Daw et al. (2006); Pearson et al. (2014)), information seeking (Geana et al. (2016)), and cognitive flexibility (Sandbrink and Summerfield (2024)). Some analyses of participant performance in this dataset have been reported previously (Knep et al. (2024); Yan et al. (2024); Mendelson et al. (2023)) and this data has been used in model development for machine learning (Mendelson et al. (2023)), but all methods and analyses described here are new.

**Figure 1:**
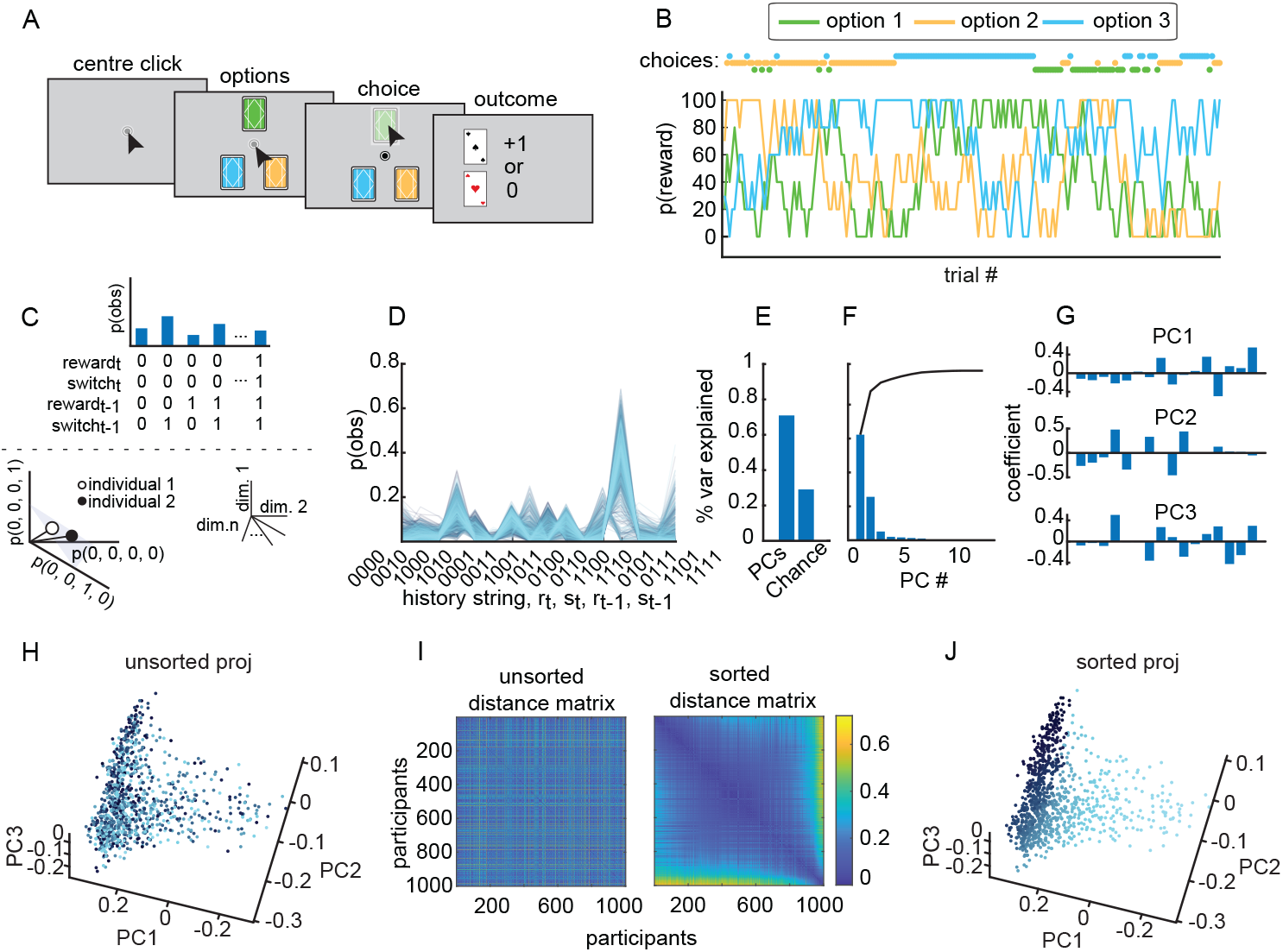
Experiment paradigm and manifold position. *A)* Trial sequence in the restless three-armed bandit task. Participants select an option, each with fluctuating reward probabilities, and receive feedback. *B)* Reward probability dynamics for the three options, with participant choices shown above. *C)* Probability of 16 choice-reward states across trials (top) and projection in highdimensional space for two participants (bottom). *D)* Distribution of the 16 choice-reward states across participants. *E)* Explained variance of principal components (∼ 71%) vs. chance (∼ 29%), computed using confound-null PCA (cnPCA). *F)* Explained variance of the 16 states across principal components (PCs). *G)* Coefficients of the 16 states for the first three PCs (cnPCA), same order as D. *H)* 3D projection of participants in PC space, with colors showing participant order. *I)* Heatmap of Euclidean distances between participants in the first two PCs (left: unsorted, right: sorted using multidimensional scaling). *J)* 3D representation of participants in top three PCs, sorted via multidimensional scaling. Dark blue-to-light blue gradient shows increasing distances in high-dimensional space.

### 3.1 Individual differences in the explore/exploit task lie on a low dimensional manifold

To characterize the ways in which individuals differed in this task, we (1) re-represented each individual’s unique pattern of choices and rewards as a point in a high dimensional space, and then (2) used dimensionality reduction to identify and rank individuals along the principle axis of variation (see Methods 5.4 and 5.5). Unlike methods that rely on cognitive modeling (Sandbrink and Summerfield (2024)) or behavioral measures, such as win-stay probabilities (Worthy et al. (2013)), this approach allowed us to determine how people differed without making strong assumptions about their approach to the task, the computations they used, or the number of ways in which they would differ from each other.

Because reward walks were stochastic, we developed a novel dimensionality reduction method to orthogonalize behavior from this stochastic aspect the task (confound-null Principle Components Analysis [cnPCA]; see Methods 5.4). Using cnPCA, we found that the participants differed along a small number of linear dimensions. Together, the first three dimensions explained 70.93% of the variance between individuals that was not due to differences in the task (Fig. 1E; differences in chance reward probability explained 29.07% of the variance). The factor loadings of the first three principle components are illustrated in Fig. 1G.

Visual inspection of the cloud of points or “manifold” of individuals (Bieleke et al. (2024)) in the cnPCA space (Fig. 1H) suggested that the majority of interindividual variance in this task was confined to a single principle dimension that curved through the space. In order to estimate each respondent’s position with respect to this nonlinear principle axis, we applied a simple sorting algorithm based on multidimensional scaling (5.5). The algorithm generated individual labels that largely corresponded to position along the principle nonlinear axis that was apparent in the cnPCA space (Fig. 1J). Hereafter, we will refer to each individual’s position along the principle nonlinear axis of the manifold of individuals as their “manifold position”.

### 3.2 Manifold position strongly and selectively co-varies with exploration

In order to determine what factors drove inter-individual variability in this task, we next asked which task variables, if any, were most strongly aligned with manifold position. First, as expected, we found that manifold position was orthogonal to chance reward probability (Fig. 2A, inset; r = −0.02, p = 0.46, Pearson correlation). This validated the efficacy of the cnPCA approach in controlling for nuisance effects of stochasticity in the reward walks. However, there was still a significant correlation between manifold position and the reward each participant earned in the task (Fig. 2A). This suggests that the principle dimension of differences did align with task performance, above and beyond what was expected by chance.

**Figure 2:**
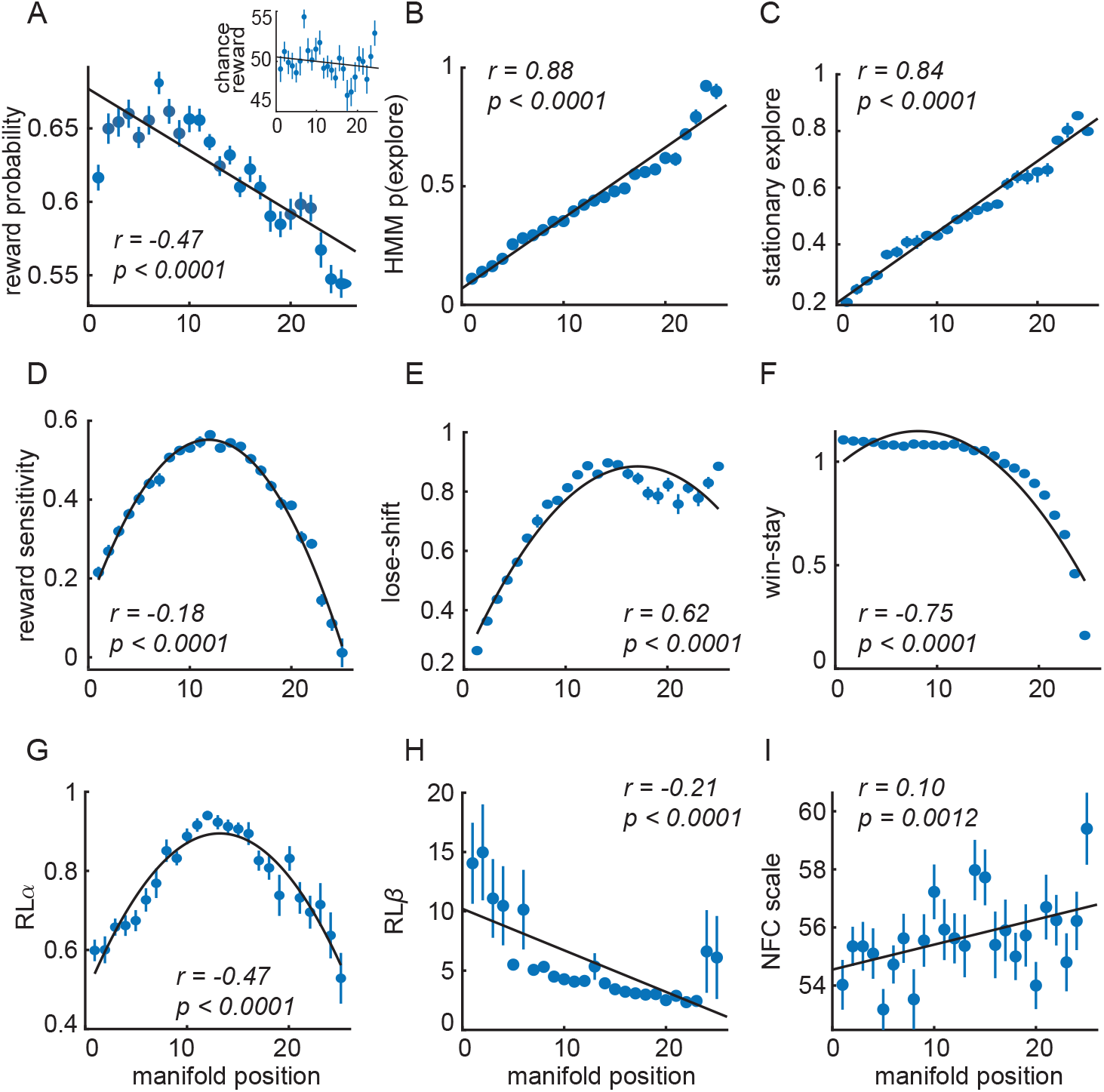
Exploration manifold position and bandit task performance. *A)* Correlation between manifold position and received reward in the main panel, and chance reward in the inset. The probability of receiving a reward (y-axis), plotted as a function of quantile-binned manifold position (x-axis; 25 bins). Each dot represents the mean received reward for each manifold position. Error bars = standard error of the mean (SEM) across participants (N = 1001). The black line represents the linear fit of the data. The legend indicates the Pearson correlation coefficient (r) and statistical significance (p). The inset is the same as the main panel but for chance reward. *B-I)* Same as (A), except the y-axes represent, in order, the HMM explore probability, stationary exploration, reward sensitivity, lose-shift, win-stay, RL *α*, RL *β*, and NFC scale. For reward sensitivity, lose-shift, win-stay, and RL *α* (panels D–G), a second-degree polynomial fit is illustrated instead of a linear fit, though the reported statistics are from a linear fit.

To determine which latent cognitive factors or task strategies influenced manifold position, we considered a range of options. These included certain model free analyses, like the probability of switching to another option following an omitted reward (lose-switch), the probability of staying on the current option following reward (win-stay), and a general reward sensitivity index (see Methods 5.6). They also included several factors that can be extracted by fitting computational models to the data, including the probability of exploration that is estimated using a Hidden Markov Model (see Methods 5.8) and the parameters of a simple Rescorla-Wagner reinforcement learning (RL) model (*α* and *β*; see Methods 5.7). We included one personality scale: the ‘need for cognition’ (NFC) scale (Cacioppo and Petty (1982)). The NFC was chosen because it is one of the stronger personality-level predictors of individual differences in curiosity and information-seeking across multiple tasks (Jach et al. (2023)).

All examined factors were significantly correlated with manifold position (Fig. 2B-I; all p *<* 0.002), consistent with previous studies linking each of these variables to individual differences in task performance. However, the single strongest predictor of manifold position was the probability of exploration, as inferred by the labels of the hidden Markov model (Fig. 2B). This was followed closely by another HMM-derived measure: the stationary probability of exploration as inferred from the fitted transition matrices (Fig. 2C). Intriguingly, manifold position was correlated with reward sensitivity, but this relationship was obviously nonlinear (Fig. 2D). One interpretation of this result is that only participants who explored at more intermediate levels were sensitive to reward: participants who either explored constantly or not at all effectively ignored the rewards. The two model-free analyses showed notable correlations with manifold position: lose-shift (Fig. 2E) and win-stay (Fig. 2F). RL model parameters (*α* and *β*, see Methods 5.7) were also correlated with manifold position (Fig. 2G-H). RL *α*, like reward sensitivity, had the same inverted U-shaped relationship suggestive of floor effects in participants who explored either constantly or extremely rarely. The NFC scale exhibited the weakest correlation with manifold position, thought this was still significant correlation (Fig. 2I), indicating a modest relationship between individual differences in cognitive style and task performance. Together, these findings suggest that factors related to exploration, reward-sensitivity, and personality each predict some of the diversity of participant behavior in the task.

In order to determine which factor, if any, made the largest contribution to variance we observed between participants, we directly compared the variance explained by each factor (see Methods 5.11; Fig. 3A). Results from both a forward selection (Fig. 3A, top) and backwards elimination (Fig. 3A, bottom) model selection procedures were consistent. The probability of exploration accounted for the largest proportion of variance in manifold position (*R*^2^ = 0.77), followed by win-stay (*R*^2^ = 0.56). Eliminating the probability of exploration from a full model that included all terms produced the largest drop in *R*^2^ (Δ*R*^2^ = 5.5%), where dropping the win-stay term was the next most effective (Δ*R*^2^ = 2.1%). It is important to note that some of these predictors were collinear (Fig. 3B), with HMM p(explore) being strongly correlated with the two nextmost-important predictors: win-stay and lose-shift. RL *α* and reward sensitivity were also correlated with win-stay and lose-shift, but neither reward-learning predictor had any appreciable ability to explain variance in manifold position in either the forward selection procedure (both *R*^2^ < 0.06) or backwards elimination procedures (both Δ*R*^2^ < 0.2%). These results suggest that individual differences in the probability of exploration were the single most important variable in explaining individual differences in task performance, outperforming factors related to reward learning, decision noise, and personality.

**Figure 3:**
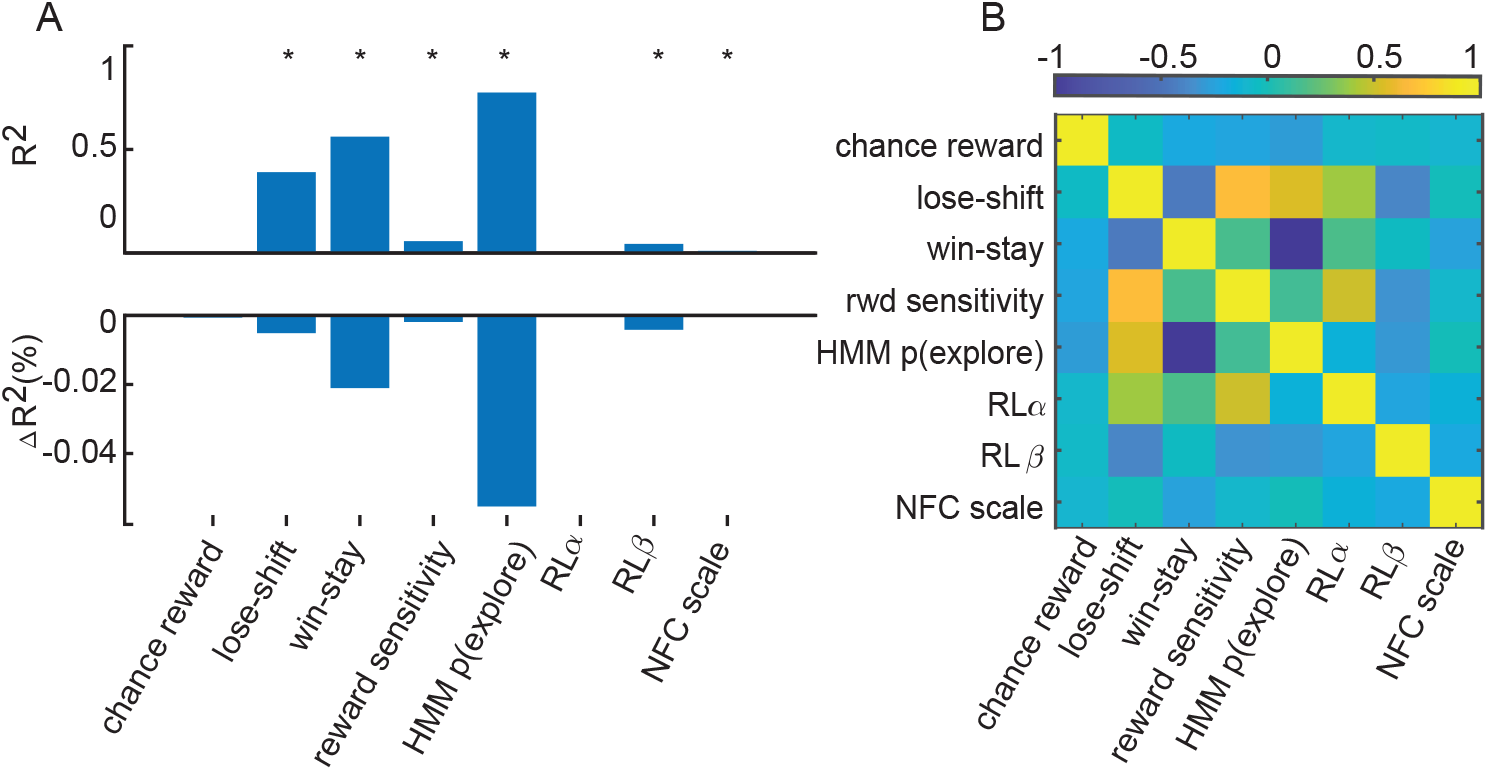
Manifold position predictors and their pairwise correlations. *A)* The top panel compares *R*^2^ values across all factors presented in panels 2A–I. The bottom panel shows the change in *R*^2^ when each factor is removed from the model. *B)* Pairwise correlation matrix of all task performance factors. The color bar represents correlation strength (Pearson*ρ*).

### 3.3 Manifold position predicted the complexity and predictability of task strategy

We next considered the possibility that individual differences in manifold position were due to differences in the randomness or complexity of the participants’ strategy in the task rather than exploration *per se*. To do this, we calculated the entropy and complexity of participants’ strategies (see Methods 5.12). First, we calculated the entropy of the distribution of choice-reward states illustrated in Fig. 1C-D. This is a measure of the unpredictability in a participant’s pattern of choices and rewards (Fig. 4A, top). A participant who is essentially unstructured, either because they are frequently changing their strategy or because they are behaving randomly, will have a highly entropic distribution of choice-reward states. We found that the participants with higher manifold positions (e.g. more exploration) tended to also have more entropy in their choice-reward state distributions (Fig. 4A, bottom). The fact that this relationship was highly non-linear, however, suggested that differences in choice-reward state entropy were not the major driver of individual variability.

**Figure 4:**
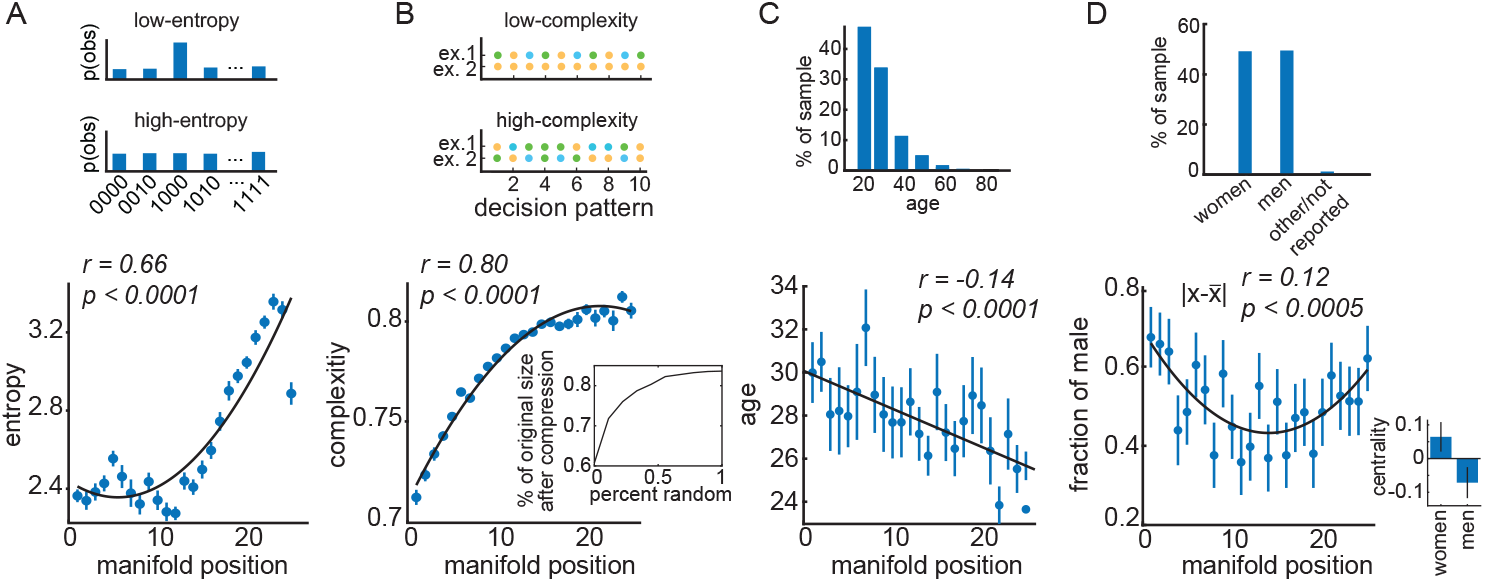
Manifold position, decision strategies and demographic factors. *A)* Top: Examples of low- and high-entropy reward and choice distributions. Bottom: Entropy plotted as a function of manifold position quantile bin. Bars are SEM and the black line is the second-degree polynomial fit. The legend indicates the Pearson correlation coefficient (r) and statistical significance (p). *B)* Top: Examples of low- and high-complexity choice sequences. Each color corresponds to a card choice (see Fig.1A-B), with changes in color reflecting the choice to switch rather than stay. Bottom: same as A, with complexity plotted as a function of manifold position. Inset: Simulated data showing how increasing the unpredictability (here, randomness) of a choice sequence increases complexity. *C-D)* Top: The percent of participants in each age bin and gender group, respectively. Bottom: Age (C) and gender (D) plotted as a function of manifold position quantile.

Next, we calculated the complexity of the participants’ strategies by measuring the compressibility of their choice sequences. We operationalized complexity as the redundant information in the choice sequences: the extent to which a file containing the choice sequences can be compressed (see Methods 5.12; Pelz and Kidd (2020)). Choice sequences that have many repeating motifs (like 1, 2, 3 or repeating 1’s) can be substantially compressed, whereas random or highly complex sequences of choices cannot be compressed to the same extent (Fig. 4B, top and inset). There was a strong positive correlation between manifold position and complexity, suggesting that participants who exhibited less structured or more complex strategies were ranked higher along the principal axis of variation.

These findings suggest that manifold position, though largely determined by the frequency of exploration, was also aligned with the complexity and predictability of decision-making strategies.

### 3.4 Age and gender differ across the manifold

To determine whether real-world personal characteristics played a role determining manifold position, we next asked whether there was a relationship between manifold position and self-reported demographic information (age and gender, Fig. 4C and D, respectively). We found that age was significantly correlated with manifold position. This implies that the age of the participant was one of the major demographic factors driving individual variability in this task.

Although self-reported gender did not have a significant linear correlation with manifold position (Fig. 4D; r = −0.04, p = 0.18), there was a moderate U-shaped relationship between manifold position and gender: gender predicted deviation from the center of manifold position (r = 0.12, p *<* 0.0005). This result is consistent with models that suggest that males are more cognitively variable than females (e.g. Arden and Plomin (2006)) and inconsistent with models that posit greater variability in females because of the uncontrolled influence of estrus cycling (e.g. Miller and Halpern (2014)). In order to determine whether this was true in all dimensions (versus just along the principle axis of the manifold) we calculated each participants “centrality” as the inverse of their median distance to every other participant in the full cnPCA space (zscored). Here again, we found that female participants tended to be more central than male participants even in the full space (p *<* 0.04, t(987) = 2.14). While other/non-reporting participants tended to be even more central, these 12 participants were not significantly different from the rest of the population (average centrality = 0.31 *±* 1.05 STD, compare male = −0.07 *±* 1.02, female = 0.06 *±* 0.97; p = 0.28, t(999) = −1.08). Together, these results suggest that the manifold of individual performance in this bandit task can be mapped back on to salient demographic variables, in addition to capturing important differences in task strategy.

### 3.5 Exploration decays exponentially with age

Given that age was a major demographic determinant of manifold position, manifold position was strongly related to exploration, and previous studies have found changes in cognitive flexibility with age (Berry et al. (2016); Kupis et al. (2021); Richard’s et al. (2021); Wilson et al. (2018); Xia et al. (2024); Mizell et al. (2024); Phelps et al. (2025)), we next asked how exploratory behavior changed with age here. First, we found that age did not predict task performance, as measured by the percent reward received minus chance (Fig. 5A). Nonetheless, despite essentially identical performance, older participants were less exploratory. Both the Hidden Markov Model probability of exploration (Fig. 5B) and the stationary probability exploration (Fig. 5C) decreased with age, indicating a shift toward more predictable and exploitative decision-making strategies in older adults. Intriguingly, in both measures of exploration, we found that a model that assumed that exploration decayed exponentially with age was a better fit to the data than models in which it changed either linearly or quadratically (see Methods 5.13; HMM p(explore), Bayesian Information Criterion [BIC] values: linear = 60,537, quadratic = 60,536, exponential decay = 59,095, all alternative BIC weights *<* 0.0001; stationary explore, BIC: linear = 54,588, quadratic = 54,584, exponential decay = 53,974, all alternative BIC weights *<* 0.0001). In short, older adults were able to achieve equivalent performance to younger adults, even as they adopted a more exploitative strategy in this flexible decision-making task.

**Figure 5:**
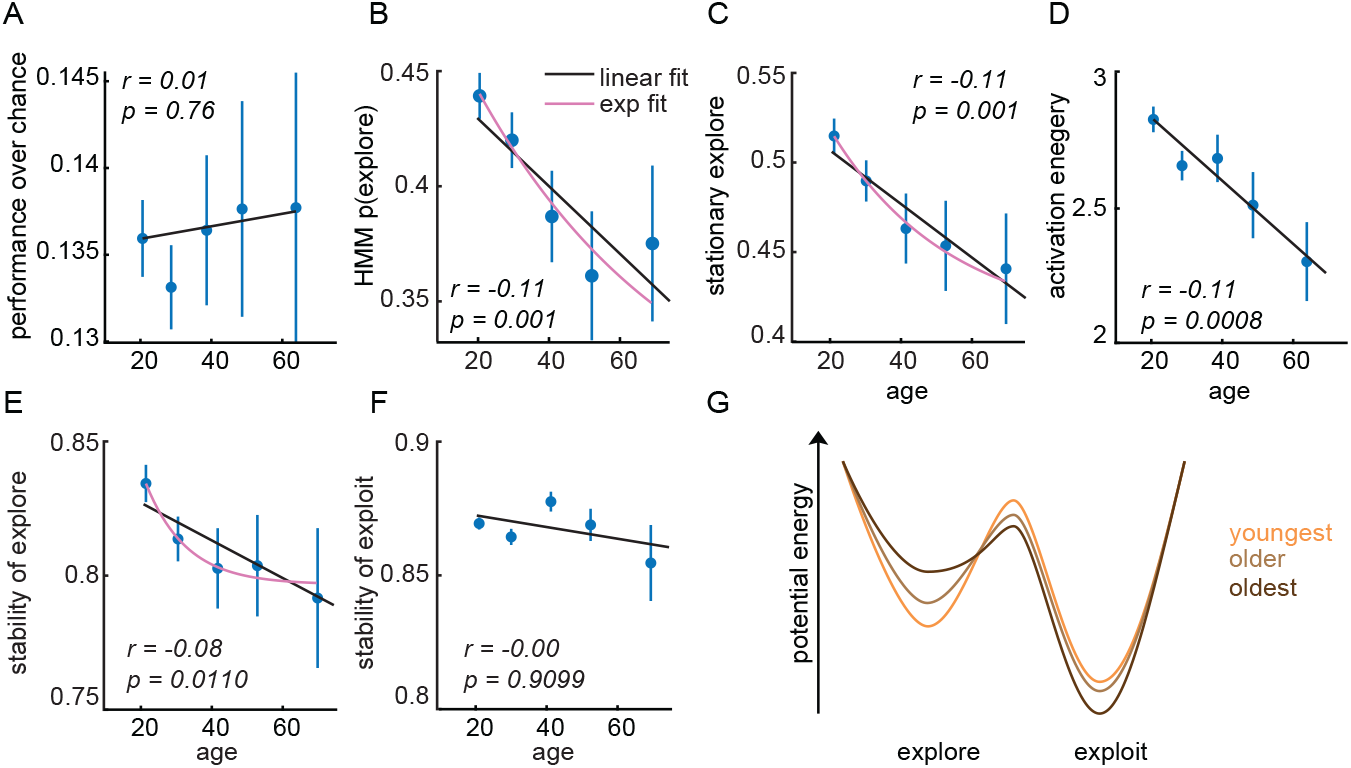
Age and performance in bandit task. *A-F)* Changes in task performance and exploration as a function of age of performance over chance (A), the probability of exploration (B), stationary exploration (C), activation energy (D), the probability of repeating an explore choice (E), and the probability of staying in an exploit state (F). The x-axis represents age bins. Black lines = linear fits, pink lines = decaying exponential fits. Insets: Pearson correlation coefficient (r) and significance (p). *G)* The energetic landscape of exploration and exploitation, illustrating differences in state dynamics between the youngest, older, and oldest participant groups, tercile binned.

In order to understand the decay in exploratory behavior in older adults, we next examined the dynamics of the HMM. Fitting the HMM involves identifying the parameters of equations that describe the transitions between exploration and exploitation in each participant (see Methods 5.9). This means that we can perform certain mathematical analyses of these equations in order to generate insight into how the dynamics of exploration and exploitation change with age (Fig. 5D-F). First, we found that the energy barrier between exploration and exploitation (i.e. “activation energy”) flattened with age: older adults had less separation between the two strategy states, suggesting more frequent transitions between them (Fig. 5D). By contrast, we found that there was no systematic relationship between age and the stability of exploitation, as measured by the probability of staying in this state (Fig. 5F). There was, however, a correlation between age and the stability of exploration (Fig. 5E). Here again, we saw evidence of exponential decay with age (BIC values: linear = 43,027, quadratic = 43,005, exponential decay = 42,971, all alternative BIC weights *<* 0.0001), suggesting that the exponential decay in the probability of exploration with age could have been due to exponential decay in the stability of exploration. To provide an intuitive summary of these results, we illustrate the energy landscape of exploration and exploitation across three age bins in Fig. 5G. Together, these results suggest that older adults’ increased tendency to exploit is primarily due to age-related decay in the stability or persistence of exploratory states.

## 4 Discussion

We found that individual differences in a restless three-armed bandit task were structured along a nonlinear low-dimensional manifold, with the principal axis of variation primarily reflecting differences in the tendency to explore. We identified this manifold via a novel dimensionality reduction method (cnPCA), which we show effectively controlled for stochasticity in the task. Because the resulting manifold was obviously nonlinear, we used a nonlinear method (multidimensional scaling) to sort people along the primary axis of the manifold. The fact that manifold position corresponded to the probability of exploration, as inferred from a HMM, was striking for three reasons. First, while k-armed bandit tasks are often touted as testbeds for examining exploration and exploitation (Berry and Fristedt (1985); Audibert et al. (2009); Macready and Wolpert (2002); Steyvers et al. (2009)), we show empirically that differences in explore/exploit behavior are the primary factor driving individual differences in this task. Second, this result highlights the value of latent state models, like the HMM, for measuring exploration and exploitation. Some prior studies have defined exploration differently (i.e. as errors of reward maximization (Daw et al. (2006)) or as synonymous with the noise parameter of a reinforcement learning model (Gershman (2018); Findling et al. (2019)). However, this result suggests that the HMM approach not only better explains variability within participants (Ebitz et al. (2018); Chen et al. (2021b)), but also between them. Third, these results highlight the fact that people differ, primarily, in how much time they spend exploring and exploiting. It is true that manifold position was aligned lower-level descriptors of exploratory behavior–like the entropy and complexity of choice sequences as a whole–but neither of these measures explained behavior as well as simple measures of the time spent exploring (versus exploiting).

The k-armed bandit and related tasks are often used to measure reward learning, generally by estimating the parameters of reinforcement learning models. This is important because differences in reward learning are a major feature of multiple neurobiological conditions (Chen and Vinogradov (2024); Aylward et al. (2019); Gilmour et al. (2024); Kaske et al. (2023); Knep et al. (2024); Addicott et al. (2013, 2021); Harlé et al. (2015); Dubois and Hauser (2022); Thinzar (2024); Daumas et al. (2023)). It is thus worth discussing the implications of our observation that differences in reward learning were not a major driver of individual differences here. While both reward learning and the *α* parameter of a RL model had a U-shaped relationship with manifold position, this did not suggest that either parameter was a major driver of individual differences. If it were, we would expect the variance between individuals to increase along this axis: making a 2 dimensional manifold, rather than a curvilinear one. Instead, the most parsimonious view is that this was a byproduct of the relationship between exploration and the manifold. Participants at the extremes—people who always explored or always exploited—would necessarily be insensitive to reward because both of these absolute strategies are reward-insensitive. Of course, it is not clear whether similar patterns would appear in other reward-learning paradigms (Abbaszadeh et al. (2023)). If so, then it raises the intriguing possibility that individual variability in reward learning could arise, in part, from differences in the amount of time individuals spend employing different decision-making strategies. Indeed, previous work suggests that there can be profound differences in the rate of reward learning between explore and exploit states (Ebitz et al. (2018); Chen et al. (2021b)). Future studies are needed to determine how individual differences in strategy use interacts with reward sensitivity to shape behavior in both healthy participants and clinical populations.

We also found that participant demographics changed systematically across the manifold. In the case of age, age-related differences in exploration drove these systematic differences in manifold position. Specifically, we found that the tendency to explore–and the stability of exploratory states–declined exponentially with age. This result resonates with the idea that younger individuals tend to prioritize exploration, while older adults are less likely to seek information, investigate novel opportunities, and sample alternative environments Sumner et al. (2019); Mizell et al. (2024); Spreng and Turner (2021); Mata et al. (2013, 2009); Chin et al. (2015). Our results extend these prior studies in two major ways. First, several of these prior studies focused on the exploration options or environments that were uncertain because of their relative novelty. Considering that age seems to increase neophobia in multiple species (Sherratt and Morand-Ferron (2018)), our results provide an important demonstration that there is a decline in exploration in older adults even in the absence of novelty. Second, we show that age-related changes in exploratory behavior can be detected even in simple analyses of the dynamics of switching in an exploratory decision-making task. This creates an intriguing foundation for future work on the mechanisms of cognitive aging because these switching dynamics are sensitive to manipulations of the same catecholamine systems that are implicated in age-related cognitive decline (Ebitz et al. (2019); Chen et al. (2023); Arnsten and Goldman-Rakic (1985)).

The decay in exploration we report in adulthood is an interesting compliment to the large body of evidence suggesting that exploration, information seeking, and cognitive flexibility decrease across development Meder et al. (2021); Harms et al. (2024); Kim and Carlson (2024); Sumner et al. (2019); Sherratt and Morand-Ferron (2018); Liquin and Gopnik (2022); Gopnik (2020); Gopnik et al. (2017). Multiple compelling hypotheses could explain these trends, including the idea that information seeking may be less beneficial to individuals who have already learned more (Sherratt and Morand-Ferron (2018); Spreng and Turner (2021)) or that our social systems offload the costs of innovation onto younger members (Gopnik (2020); Gopnik et al. (2017)). Our results are part of a body of evidence that now suggests that the decline in exploration persists into adulthood (Berry et al. (2016); Kupis et al. (2021); Richard’s et al. (2021); Wilson et al. (2018); Xia et al. (2024); Mizell et al. (2024); Phelps et al. (2025)). However, an open question remains: does this trend reflect a universal shift with age, or are there meaningful individual differences? Recent work by Mizell et al. (2024) suggests the latter. While some older participants showed minimal information-seeking, others maintained exploratory behaviors at levels comparable to younger adults. Additionally, Sumner et al. (2019) found that when adults were explicitly instructed to learn about their environment, their performance matched that of children. These findings suggest that age-related changes in exploration likely exist on a spectrum. Future research with balanced age distributions and longitudinal designs will be essential to disentangle the effects of cognitive aging, life experience, and personality across the lifespan. The other demographic factor we considered—gender—did not have a simple linear relationship with manifold position. It did, however, predict the variability between participants. There were more self-identified male participants at both extremes of the decision-making manifold and male participants also tended to simply have more unusual patterns of choice. These results align with the male variability hypothesis, which posits greater cognitive variability among males compared to females (Arden and Plomin (2006)), and with previous studies showing greater male variability in decision-making strategies in animal models (Chen et al. (2021a)). By contrast, our results challenge the idea that hormonal fluctuations in females lead to greater decision-making variability (Miller and Halpern (2014)). This is because more fluctuations over time would increase the variability between individuals in a cross-sectional study (Miller and Halpern (2014)). One important caveat is that gender was self-reported in our study and our sample did not include sufficient non-binary and transgender participants to meaningfully assess broader gender-related effects. We did find that the fraction of individuals who did not report a binary gender identity was in line with estimates of the prevalence of transgender identities (i.e. 1.2%) and that these individuals were, if anything, more centralized within the population. However, we were still underpowered here and future research should include more diverse gender identities, perhaps through intentional sampling within transgender and non-binary communities. Work in animal models (such as the Four Core Genotypes mouse model; Gatewood et al. (2006); Arnold and Chen (2009)) could provide valuable insights into the biological and environmental factors that underlie these effects.

Together, our findings demonstrate the potential of manifold discovery methods in studying individual differences in cognition. By identifying the principal axis of variation in task performance, we determined that exploratory tendencies, rather than reward sensitivity or reinforcement learning strategies, were the primary source of individual variability in a classic explore/exploit task. Our approach was able to both recover and reinforce relationships between certain important demographic factors and individual differences in decision-making: highlighting the necessity of considering both age-related and gender-related variability in future studies of decision-making behavior. Ultimately, the novel manifold discovery approach we develop here has incredible potential in a variety of applications. Future research could leverage similar methods to refine task structures, ensuring they more effectively capture the specific cognitive constructs of interest in a given study. Combining this approach with methods for manifold alignment could be a powerful new way to develop computational models that can account for the full heterogeneity of individual response patterns (Zid et al. (2024)) or to compare behavior across algorithms (Zid et al. (2024); Ramírez-Ruiz and Ebitz (2024)) or species (Laurie et al. (2024)). Understanding how the dimensions of a manifold map relate to clinical symptom scores could further extend the utility of these methods into computational psychiatry and cognitive assessment (Knep et al. (2024); Yan et al. (2024)). Ultimately, manifold discovery methods have incredible potential for uncovering hidden patterns of variability across cognitive tasks, advancing our understanding of the latent cognitive variables that truly shape individual differences in decision-making and other complex behaviors.

## 5 Methods

### 5.1 Data Collection

All experimental procedures were consistent with and approved by Institutional Review Board of the University of Minnesota (Study 00008486). Participants were recruited via 2 online platforms, Amazon Mechanical (mTurk) and Prolific. In both cases, geographic and other restrictions were in place to minimize the likelihood of bots and other data quality issues. Participants were compensated for their time in proportion to the number of points that they earned (generally: $0.02 cents per point) and all participants earned at least the minimum hourly compensation rate for the platform. A total of 1001 participants completed the task (493 female, 496 mean and other non-reporting).

### 5.2 Task design

In the task, participants were required to repeatedly choose between 3 options (1A). Each option was associated with a probability of reward, which changed across trials *t* and independently across options *i* according to

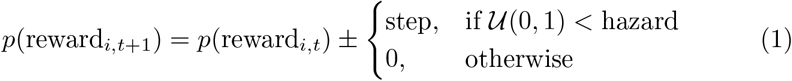

where “hazard” is a fixed rate of change ∈ [0, 1], *U* (0, 1) is a draw from a uniform random distribution, and the sign of the step is chosen independently at random for each option on each trial. Each participant’s reward schedule was randomly generated (Fig. 1B). The experiment consisted of 25 practice trials that guided the participants through the mechanics of the task, followed by 300 trials of the experiment. Only the 300 experimental trials were analyzed.

### 5.3 General analyses

Custom MATLAB scripts were used for data analysis. The relationship between manifold position and task performance factors was examined using Pearson correlation. A significance threshold of p = 0.05 was applied. Fitted lines in Fig. 2 and Fig. 4 represent linear fits, except for U-shaped trends, which were modeled using second-order polynomial fits. Because age was both reported in non-uniform bins (18-24, 25-34, 35-44, 45-54, 55-64, 65-74, 75+) and not uniformly sampled (Fig. 4D), participant ages were recoded to bin centers that corrected for this geometric fall off (i.e. people in the 18-24 bin were coded as 20.59, rather than 21, 25-34 were coded as 28.67, not 29.5, etc.). Statistics that were run on raw, recoded age data are reported in the text and eliminating this correction did not alter the results. Because there were less than 20 participants in all age bins greater than 55, these bins were combined for illustration and centered at the geometrically-corrected center of the combined bin (e.g. in Fig. 5).

### 5.4 Manifold discovery

In order to determine how individuals differed in the task without regard to any specific hypotheses about the computations they were performing, we re-represented behavior as probability distributions over strings of reward and choice history (Yan et al. (2019); Chen et al. (2021a)). For each participant, we calculated the probability of occurrence of specific strings of *n* prior rewards (coded as 1 for reward, 0 otherwise) and *m* switch choices (coded as 1 for switch, 0 otherwise), meaning that each participant’s choices were coded as a probability vector with length 2^*n*^ * 2^*m*^. Geometrically, that means that we re-represented each participant as a points on a 2^*n*^ * 2^*m*^ dimensional probability simplex (Fig. 1H-J).

Next, we performed dimensionality reduction to understand individual differences in choice patterns. Principal component analysis (PCA) is the most common method for reducing the dimensionality of high-dimensional data sets for visualization and analysis. PCA identifies a set of orthonormal ‘principal’ axes that capture covariance between data points, ordered by decreasing explained variance, without considering the underlying cause of the variance. In contrast, we aimed to distinguish individual differences in task behavior from the effects that the task itself might have on individuals. To separate these two factors, we developed a novel dimensionality reduction method that we call confound-null PCA (cnPCA). Briefly, cnPCA first identifies the principal axes in the data that are related to confounding variables and then performs PCA in the null space of the confounds. In the case of 1 confounding variable, we would find the *n ×* 1 vector *β* that predicts the *m ×* 1 confound vector **C** from the *m × n* data matrix **X**, where *C* = **X***β*. Geometrically, *β* then represents the projection of **X** that explains the most variance in the confounding variable *C*. For a set of *k* confounding variables, **C**, we solve for the *n × k* matrix **B** that satisfies **C** = **XB**. Given the set of projections of **X** that are related to **C**, we then iteratively calculate each PC_*k*_ as the one that maximizes covariance, subject to the constraint that it is orthogonal to **C** and also to PC_1:*k*−1_. Here, we applied cnPCA to control for differences in the chance probability of stochastically generated reward walks (Fig. 1E-G).

### 5.5 Estimating position along the manifold

In order to determine each individual respondent’s position with respect to the principle axis of variability, we applied a simple seriation algorithm (Behrisch et al. (2016)). We calculated the pairwise Euclidean distance between each individual in the cnPCA space, and then performed multidimensional scaling on the pairwise distance matrix (Matlab: *mdscale*, criterion = metric stress). Briefly, multidimensional scaling is a nonlinear dimensionality reduction algorithm that remaps a set of points into an abstract space where points that were close together in the original (unperturbed) space remain close together in the remapped space (Fig. 1H, before sorting; J, after sorting). By sorting individuals according to their position along the first output dimension of multidimensional scaling, we were able to assign positions to each individual that visually corresponded to their position along the nonlinear axis of co-variability that was apparent in the original cnPCA space (Fig. 1J).

### 5.6 Lose-shift, win-say and reward sensitivity

In order to look at differences in reward learning and reward sensitivity without assuming a particular learning model, we examined the probability of switching, conditioned on the previous trial’s outcome. We also used the probability of switching conditioned on previous trial reward outcome to calculate reward sensitivity using a common 1-trial-back index of reward learning,

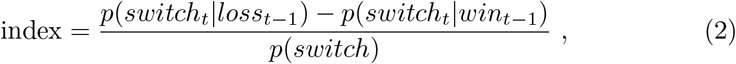

where *p*(*switch* | *outcome*) is the conditional probability of switching on one trial, given the outcome of the previous trial (Fig. 2D-F).

### 5.7 Reinforcement learning model

A reinforcement learning (RL) model was used to examine the correlation between RL parameters and manifold position. The model follows the standard Rescorla-Wagner learning rule, which updates the value *v* of chosen option *i* on each trial *t* according to the delta-rule:

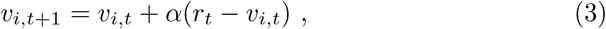

where *r*_*t*_ represents the reward on trial *t* (0 for no reward, 1 for reward) and *α* ∈ [0, 1] is the learning rate parameter of the model. The delta-rule update dynamically calculates value as an exponentially-decaying recency-weighted average of reward. The value of unsampled options is not updated, so *v*_*k,t*+1_ = *v*_*k,t*_ for all unsampled options *k*.

Values are then transformed into choice probabilities via passing them through a softmax,

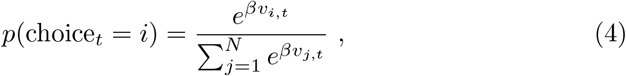

where *β* ∈ [0, 100] is an inverse temperature parameter that controls the noise in the value comparison. Models were fit to individual participants initialized via maximum likelihood (minimizing the negative of the log likelihood; Matlab: *fminsearch*) and fits were reinitialized with 20 random seeds.

### 5.8 Hidden Markov model

To identify when both models and participants were engaging in exploration versus exploitation, we implemented a hidden Markov model (Ebitz et al. (2018, 2020); Chen et al. (2021b)). In this framework, choices (*y*) are considered ‘emissions’ that resulted from an unobserved decision-making process, which operates in a latent, hidden state (*z*). These latent states are characterized by both the probability of selecting a particular choice (*k*, out of *N*_*k*_ possible options), and by the probability of transitioning between states. Our model included two distinct types of states: an explore and multiple exploit states. The emission model for the explore state followed a maximum-entropy distribution for a categorical variable, which corresponds to a uniform distribution:

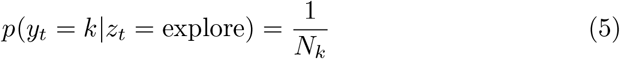

We aimed to minimize the number of assumptions made about the choices during exploration to avoid introducing any bias that could favor or discourage a particular policy type. However, using a uniform distribution to model exploratory choices does not suggest, necessitate, or impose random decisionmaking during these states(Ebitz et al. (2020, Ebitz et al. 2019)). On the other hand, because exploitation involves repeatedly selecting the same option, exploit states only allowed choice emissions corresponding to a single option. That is:

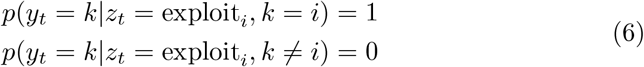

The latent states in this model follow a Markovian process, meaning that the current state (*z*_*t*_) is determined solely by the previous state (*z*_*t*−1_):

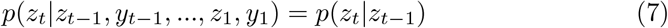

This implies that the entire pattern of dynamics can be represented by a time-invariant transition matrix between four possible states: three exploit states and one explore state. This matrix consists of stochastic equations that define the probability of transitioning between all possible combinations of past and future states (i, j) at each time step.

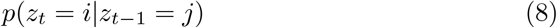

Due to the limited number of trials per participant (300), parameters were shared across exploit states: each exploit state had the same probability of initiating (from exploration) and of maintaining itself. For conceptual clarity, the model also assumed that participants began in the exploration state and needed to pass through it before transitioning to exploiting a new option, even if it was only for a single trial (Ebitz et al. (2018, 2020); Chen et al. (2021b)). We have previously demonstrated that models without these design constraints tend to approximate them when fitted to sufficiently large datasets (Chen et al. (2021b); Ebitz et al. (2018)).

Since the emissions model was fixed, certain parameters were shared, the structure of the transition matrix was restricted, and the initial state was predetermined, the final HMM had only two free parameters: one representing the probability of exploring given exploration on the previous trial, and the other representing the probability of exploiting given exploitation on the previous trial. We have previously shown that this constrained model performs equally well as an unconstrained model (Chen et al. (2021b); Ebitz et al. (2018)) and that unconstrained models tend to closely align with the constrained model when fitted to large datasets (Chen et al. (2021b); Ebitz et al. (2018)).

The HMM was fitted using expectation-maximization with the Baum-Welch algorithm (Bilmes (1998)). This algorithm identifies a (potentially local) maximum of the complete-data likelihood. The algorithm was restarted with random seeds 20 times, and the model that maximized the observed (incomplete) data log likelihood across all sessions for each participant was selected as the best. To infer latent states from choices, we applied the Viterbi algorithm to determine the most probable a posteriori sequence of latent states.

### 5.9 Analyzing HMM dynamics

In order to understand how the balance between exploration and exploitation changes across manifold position, we analyzed the “dynamics” of the HMM. We used tools from statistical thermodynamics to characterize the long-term behavior and potential energy landscape of the fitted latent state models.

In statistical mechanics, the energy of a state (like exploration or exploitation) is related to the long-term probability of observing a process (like a decision-maker) in those states. A low-energy state is one that is very stable and deep and will be over-represented in the system’s long-term behavior. A high energy state is less stable and will be under-represented in the system’s behavior. The Boltzman distribution relates the probability of observing a process in a given state *s* (*p*_*s*_) to the energy of that state (*E*_*s*_),

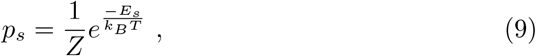

where *Z* is the partition function of the system, *k*_*B*_ is the Boltzman constant, and *T* is the temperature. If we consider the ratio between two states, *s* and *k*, the partition functions cancel out and the relative occupancy of the two states is just a function of the difference in free energy between them,

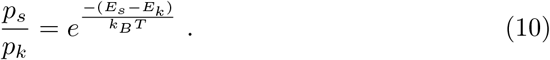

Rearranging, we can express the difference in energy between the two states as a function of the difference in the long-term probability of those states being occupied,

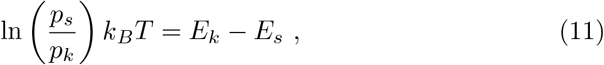

meaning that the difference in the energetic depth of the states is proportional to the log odds of each state, up to some multiplicative factor kBT.

Because of how parameters were tied in the HMM, the model effectively had two states: an explore state and a generic exploit that described the dynamics of all exploit states. We estimated the probability of exploration and exploitation for each subject *i* by calculating the stationary distribution of the HMM’s transition matrix for subject *q* (*A*_*q*_). That is, we found the probability distribution *π*^*^ that satisfied

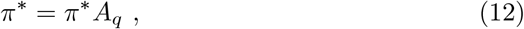

meaning the probability distribution over states that remained unchanged by repeated application of the fitted dynamics. If the transition matrix is ergodic, the stationary distribution is the (normalized) left eigenvector of the transition matrix with an eigenvalue of 1. A small number of participants were excluded because HMM transition matrices did not admit a stationary distribution (3.7%, 38/1001). The term “stationary explore” is used to refer to the difference in the stationary distribution between explore and exploit states.

In order to understand the dynamics of exploration and exploitation, we also need to estimate the energy barrier between the states. Here, we use the Arrhenius equation from chemical kinetics to relate the rate of transitions, *r*, between some pair of states to the activation energy required to affect these transitions, *E*_*a*_,

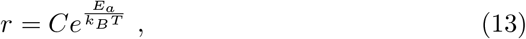

where *k*_*B*_*T* is the product of temperature and the Boltzman constant and *C* is a constant pre-exponential factor. Rearranging, we can solve for activation energy,

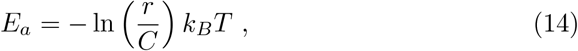

where *r* is the rate of transitions given by the entries of the HMM transition matrix. *E*_*a*_ was calculated bidirectionally (i.e. from *r*_*ore*→*oit*_ and *r*_*oit*→*ore*_). Where different solutions were found (difference *>* 10^−4^), participants were excluded from additional analyses. Note that these equations can only identify three discrete points on the energy landscape (an explore state, an exploit state, and the peak of the barrier between them). These 3 points were calculated within participants then averaged within participant groups. A continuous potential was traced through the means to provide a physical intuition for the changing energy landscape (Fig. 5E).

Finally, we investigated the correlation between the fraction of time spent in exploration (stationary explore) for each participant and manifold position. We also examined how the HMM dynamics parameters change with age (Fig. 5B).

### 5.10 Need for cognition (NFC) scale

Need for Cognition (NFC) is a psychological construct that refers to the extent to which individuals enjoy and engage in effortful cognitive activities, including their curiosity, openness to novel information, and willingness to explore new strategies (Cacioppo and Petty (1982)). It reflects a person’s tendency to actively seek out and enjoy complex thinking, problem-solving, and the acquisition of new information (Cacioppo and Petty (1982)). To quantify NFC in the context of our task, we used the Need for Cognition Scale (NFC inventory), a self-report measure developed by Cacioppo and Petty (1982). The NFC inventory assesses individual differences in the tendency to enjoy thinking and the propensity to engage in cognitive efforts (Cacioppo and Petty (1982)).

In this context, a higher NFC score indicated a greater tendency to engage in exploratory behavior—seeking new information and options—rather than strictly exploiting the options that seemed most rewarding based on prior experience. This aligns with recent frameworks on information demand in decisionmaking (**?**), which emphasize the role of individual differences in the pursuit of new information and the process of exploration in dynamic environments.

### 5.11 Generalized linear models

In order to compare the relative contribution of multiple factors to manifold position *y*, we used generalized linear models to perform simplified forward selection and backward elimination model comparison. First, for forward selection, we fit a series of models for each individual predictor *X*_*k*_,

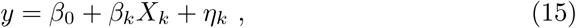

where *η*_*i*_ is the noise term of model *k* and *β*_*k*_ is the coefficient of predictor *k*. We then measured the coefficient of determination of each model 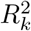,

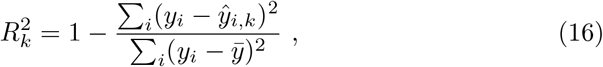

where *ŷ*_*i,k*_ is the estimate for observed manifold position *i* from model *k* and 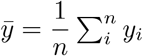. The best model was taken as the one with the largest *R*^2^.

In the backward elimination process, we began with a model that included all *N* predictors,

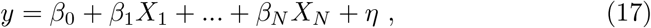

then eliminated each term, one-by-one, while calculating the percent change in the coefficient of determination in the model that omitted predictor *k*,

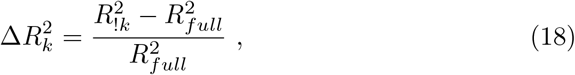

where the coefficient of determination of the full model is 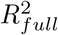 and the equivalent for the model without predictor *k* is 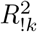.

### 5.12 Entropy and complexity

To analyze participants’ decision-making behavior, we examined two parameters: entropy and complexity. These measures capture different aspects of how participants navigated the decision space in our three-arm bandit task.

Entropy quantifies the unpredictability of participants’ choices. A higher entropy value indicates that all possible choice patterns occur with similar probabilities, suggesting a more exploratory strategy. In contrast, lower entropy suggests a preference for a limited set of decision patterns (Fig. 4A).

To compute entropy, we first calculated the probability of occurrence of specific choice sequences based on the two prior choices (switch vs. stay) and two prior reward outcomes (rewarded vs. not rewarded). We then calculated entropy for each participant using the Shannon entropy formula:

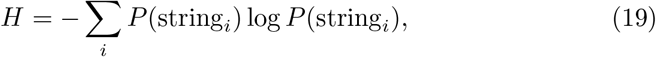

where *H* represents the entropy of a participant’s decision strategy, *P* (string_*i*_) is the probability of observing a particular choice pattern *i*, and log *P* (string_*i*_) scales the probability logarithmically.

While entropy captures the predictability of decision patterns, complexity reflects how structured or variable a participant’s sequence of choices is over trials. A highly repetitive choice pattern is considered low complexity and more compressible, whereas more varied sequences indicate greater complexity.

To quantify the complexity of participants’ decision-making strategies, we measured the compressibility of their choice sequences using ZIP compression (Pelz and Kidd (2020)). This method assesses the redundancy in decision patterns by evaluating how much the sequence size is reduced when subjected to Lempel-Ziv compression (Ziv and Lempel (1977)). Sequences that compress more efficiently indicate higher redundancy and lower complexity, whereas sequences that remain less compressed suggest greater exploration and unpredictability. For each participant, we extracted their choice sequence from the three-bandit task. The extracted sequence was then saved as an uncompressed MATLAB .mat file using version -v6, ensuring that no built-in compression was applied. Next, the file size was recorded in bytes to establish a baseline measurement. To assess compressibility, the file was then compressed using MATLAB’s zip() function, and the size of the resulting compressed file was recorded. Finally, the compression ratio was computed as the ratio of the compressed file size to the original uncompressed size:

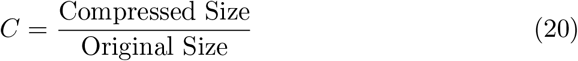

This ratio provides a measure of complexity in the decision-making sequence, where lower values indicate greater repetitiveness and lower complexity, while higher values reflect more complex and unpredictable decision patterns.

### 5.13 Decay in exploratory processes

In order to determine how exploration changed with age, we compared linear, quadratic, and exponential decay models. The models were fit to the HMM p(explore), the stationary probability of exploration, and the probability of staying in exploration. The linear model,

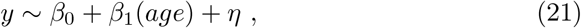

assumed a linear rate of change in these exploratory processes with age. Here, *β*_0_ and *β*_1_ represent offset and slope parameters, respectively, and *η* is the noise term to be minimized. The quadratic model,

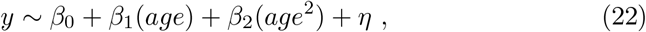

added a degree of freedom (*β*_2_) that allowed for curvature in the dependence of the parameter on age without assuming a particular functional form. By contrast, the exponential decay model,

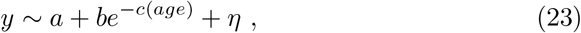

assumed that exploratory processes decayed eponentially with age. Here, age was normalized by subtracting its minimum. Because *y* ∈ [0, 1] for all *y*, likelihoods were calculated with the binomial likelihood function for each participant *i*,

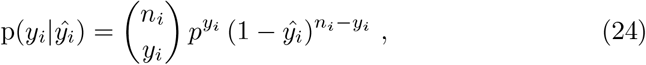

where *ŷ*_*i*_ was the predicted value of each model and *n*_*i*_ was their number of choices. Log likelihoods were then summed across participants. Bayesian information criteron values are reported in the text, but identical results were obtained using the Akaike Information Criteron.

## 5.14 Acknowledgment

This study was supported by the Jacobs Foundation (Research Fellowship), the Natural Sciences and Engineering Research Council of Canada (RGPIN2020-05577), the Canada Research Chair in the Dynamics of Cognition (CRC-2022-00192), the US Department of Defense (FA9550-24-1-0305), the National Institute of Mental Health (R21MH127607), and the National Institute on Drug Abuse (K23DA050909).

